## Supplementary for "MethylBERT: A Transformer-based model for read-level DNA methylation pattern identification and tumour deconvolution"

### Supplementary tables and figures

| | HMM/Bayesian/Random Forest-based model | CNN/RNN-based model | Transformer-based model |
|---|---|---|---|
| **Methylation site prediction** | iDNA-MS<br>SOMM4mC<br>MM-6mAPred | DeepTorrent<br>iIM-CNN | iDNA-ABT<br>MuLan-Methyl |
| **Nanopore methylation calling** | Nanopolish<br>SignalAlign | DeepMod<br>DeepSignal | Rockfish<br>methBERT |
| **Single-cell methylation imputation** | Melissa<br>CaMelia | DeepCpG | CpG Transformer |
| **Tumour purity estimation** | CancerDetector<br>BED<br>DXM<br>CAMDAC | DISMIR | **MethylBERT** |

**Supplementary Table 1.** Overview of methods for sequencing-based DNA methylome analysis

| | CancerDetector | DISMIR | DISMIR_dmr | Houseman |
|---|---|---|---|---|
| **MethylBERT** | 1.089e-09 | 2.594e-19 | 1.856e-03 | 4.013e-26 |
| **MethylBERT +adjustment** | 1.433e-09 | 1.102e-18 | 7.695e-04 | 2.392e-18 |

**Supplementary Table 2.** Paired t-test p-values with Bonferroni correction for Figure 4A. The p-value for MethylBERT vs MethylBERT+adjustment is 1.737e-04.

| | CancerDetector | DISMIR | DISMIR_dmr | Houseman |
|---|---|---|---|---|
| **MethylBERT** | 1.000e+00 | 8.828e-04 | 4.376e-09 | 1.370e-06 |
| **MethylBERT +adjustment** | 1.000e+00 | 1.228e-03 | 1.325e-08 | 1.451e-06 |

**Supplementary Table 3.** Paired t-test p-values with Bonferroni correction for Figure 5A. The p-value for MethylBERT vs MethylBERT+adjustment is 5.218e-01.

| Cell type | Samples |
|---|---|
| Prostate epithelial | Prostate-Epithelial-Z000000RV<br>Prostate-Epithelial-Z000000S3<br>Prostate-Epithelial-Z0000045F<br>Prostate-Epithelial-Z0000045G |
| Non-prostate epithelial | Blood-T-EffMem-CD4-Z00000416<br>Colon-Fibroblasts-Z0000042C<br>Blood-B-Z000000UB<br>Blood-T-CD8-Z000000U5<br>Blood-B-Mem-Z0000041J |

**Supplementary Table 4.** Normal cell-type methylation atlas samples used for training MethylBERT to estimate prostate epithelial cell-type proportions in Figures 4F and G.

| Cell type | Samples |
|---|---|
| Blood-B | Blood-B-Z000000UB<br>Blood-B-Z000000UR |
| Blood-NK | Blood-NK-Z000000UF<br>Blood-NK-Z000000U1 |
| Blood-Granul | Blood-Granulocytes-Z000000TZ<br>Blood-Granulocytes-Z000000UT |
| Blood-T | Blood-T-Eff-CD8-Z0000041Q<br>Blood-T-CenMem-CD4-Z0000041D |
| Blood-Mono+Macro | Blood-Monocytes-Z000000TP<br>Lung-Alveolar-Macrophages-Z00000448 |

**Supplementary Table 5.** Normal cell-type methylation atlas samples (Loyfer et al.) used for training MethylBERT to deconvolute leukocyte sample in Figure 4H.

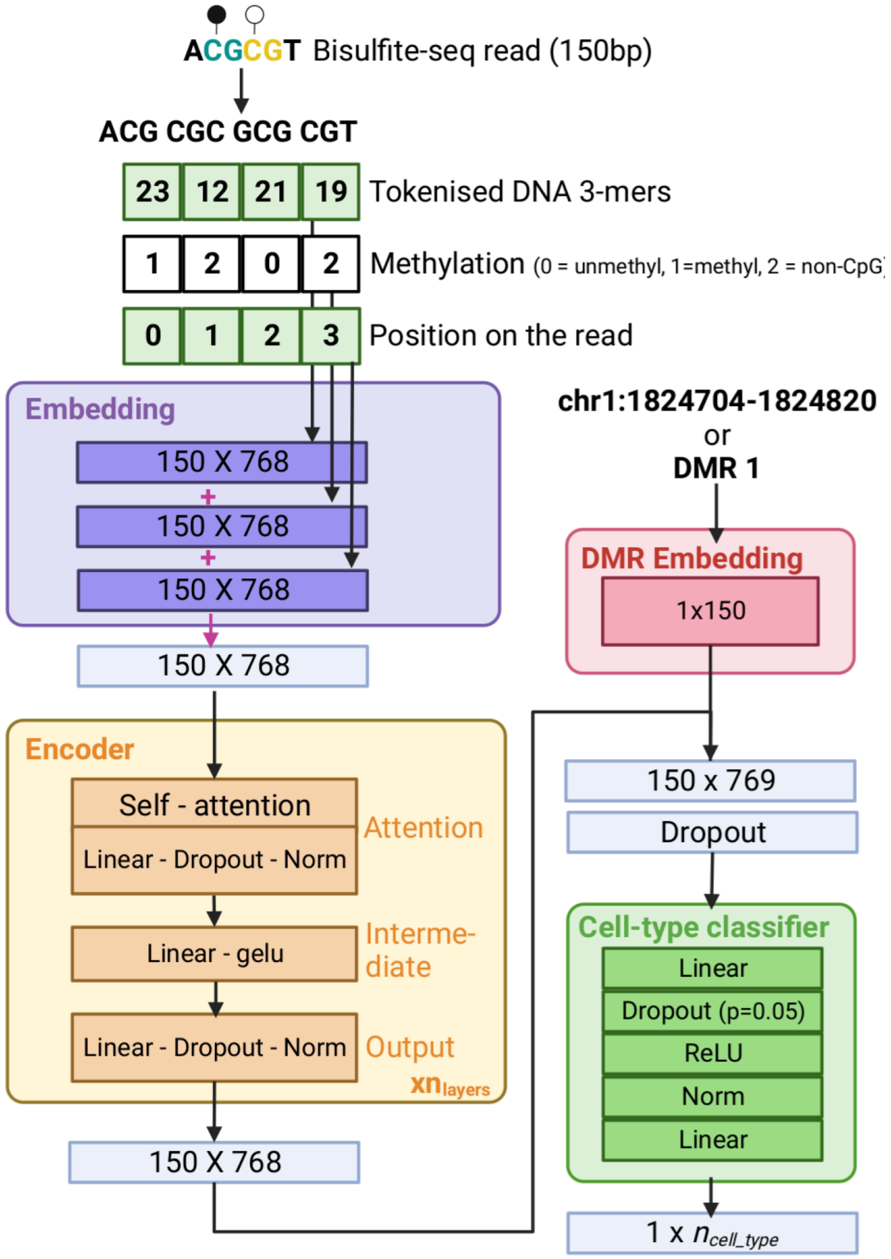


**Supplementary Figure 1.** MethylBERT network architecture

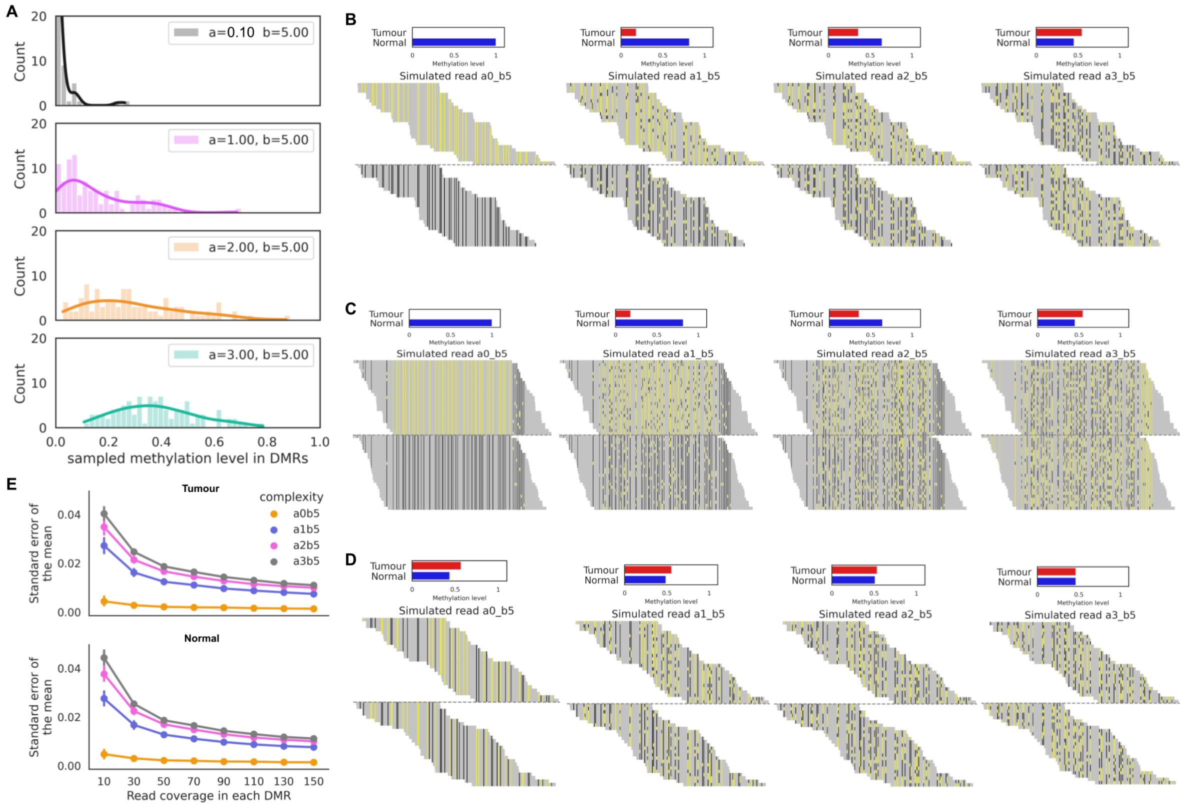


**Supplementary Figure 2.** Simulated read-level methylomes **A.** Distribution of tumour methylation level in DMRs sampled from a beta distribution with different α values. **B-D.** Examples of simulated methylation patterns in four different complexities. The read length is set up as 150bps (B) and 500 bps (C), respectively. CpG-specific methylation patterns are given instead of region-specific methylation levels (D). The grey dotted line separates tumour methylation patterns (above the line) and normal methylation patterns (below the line). The histogram at the top of the methylation patterns indicates tumour and normal methylation levels in the region. **E.** Mean-variance of methylation patterns in DMRs for different coverage of simulated data.

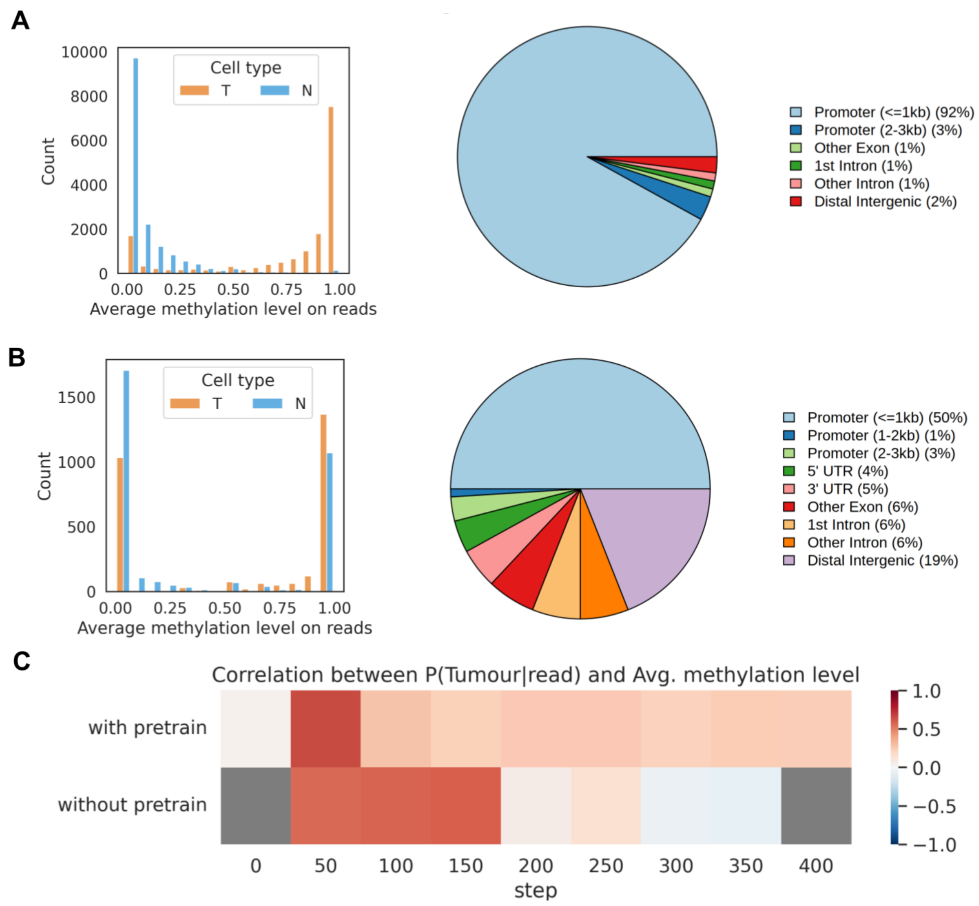


**Supplementary Figure 3.** Additional pre-training result analyses. **A.** Distribution of read-wise methylation level in DMRs selected according to areaStat score (left) and genomic annotation of DMRs (right). **B.** Distribution of read-wise methylation level in DMRs where hypomethylated and hypermethylated patterns are balanced (left) and genomic annotation of DMRs (right). **C.** Correlation between P(cell type=Tumour|read) and average methylation level in each DMR set before and after pre-training. Grey colour means an insignificant correlation value.

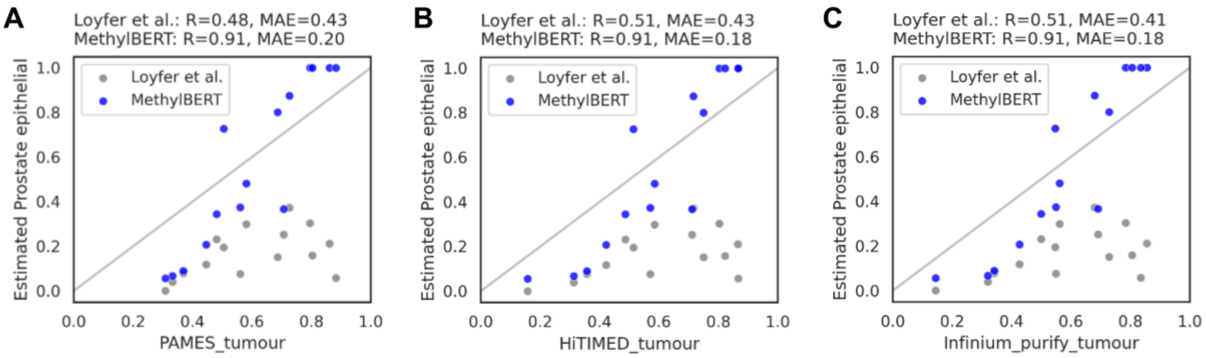


**Supplementary Figure 4.** Correlation between the estimated prostate epithelium proportion (without tumour reference data, by MethylBERT and Loyfer et al.'s method, respectively) and the estimated tumour purity using tumour reference data by PAMES (A), HiTIMED (B), and InfiniumPurify (C) for the lymph node samples acquired from hormone-sensitive metastatic prostate cancer patients. Different methods are distinguished by different colours.

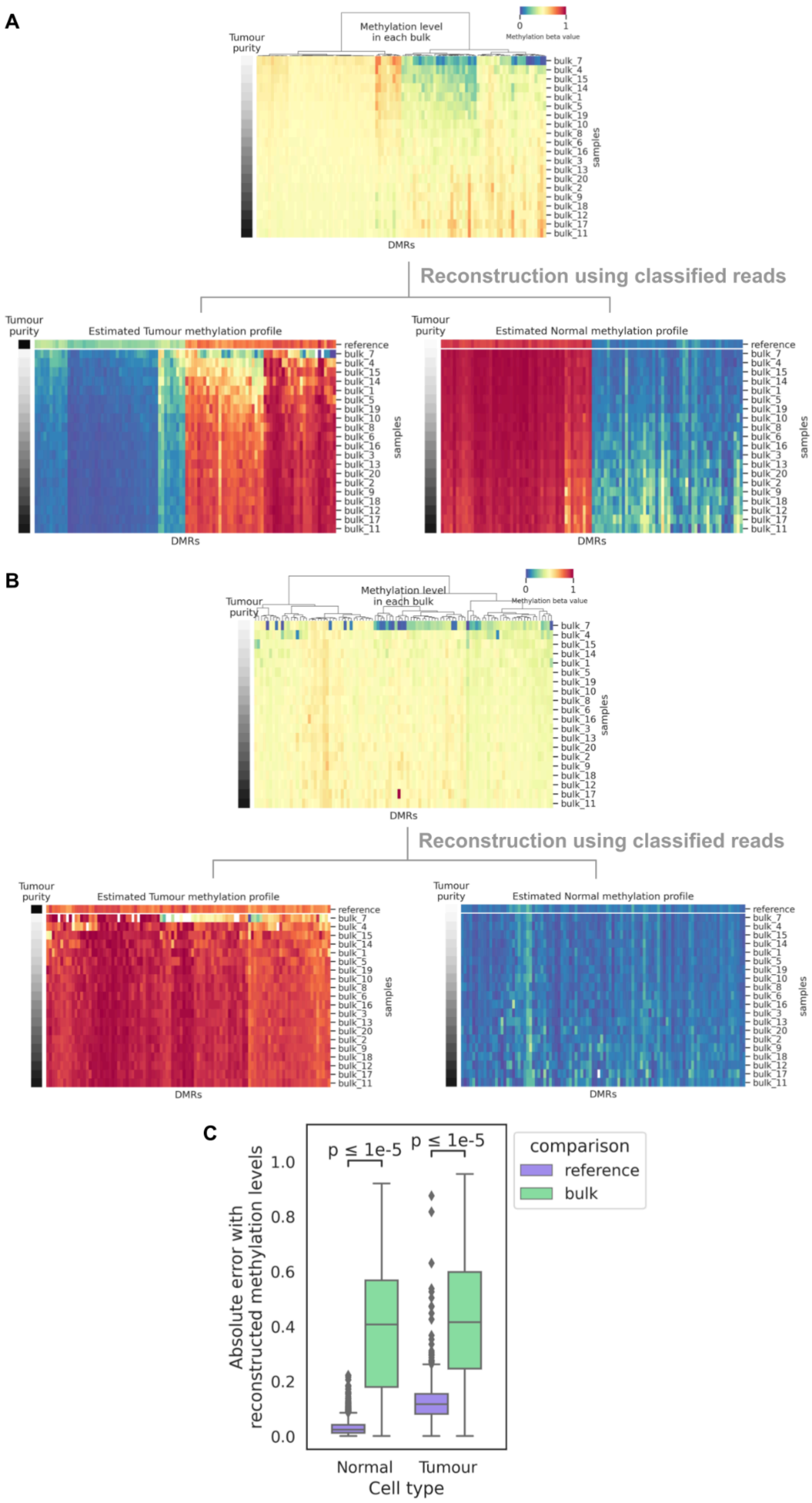
A
Methylation level in each bulk
Tumour purity
Methylation beta value
0
1
bulk_7
bulk_4
bulk_15
bulk_14
bulk_1
bulk_5
bulk_19
bulk_10
bulk_8
bulk_6
bulk_16
bulk_3
bulk_13
bulk_20
bulk_2
bulk_9
bulk_18
bulk_12
bulk_17
bulk_11
samples
DMRs
Reconstruction using classified reads
Estimated Tumour methylation profile
Estimated Normal methylation profile
reference
B
Methylation level in each bulk
Reconstruction using classified reads
Estimated Tumour methylation profile
Estimated Normal methylation profile
C
Absolute error with reconstructed methylation levels
p ≤ 1e-5
p ≤ 1e-5
1.0
0.8
0.6
0.4
0.2
0.0
Normal
Tumour
Cell type
comparison
reference
bulk

**Supplementary Figure 5.** Reconstructed DMR methylation levels in tumour and normal cell type using the MethylBERT read classification results. **A-B.** Reconstructed cell type-specific methylation patterns from all pseudo-bulks compared to reference cell type-specific methylation profiles in DMRs, when DMRs are evenly divided between tumour-hypermethylated and -hypomethylated regions (A) and are selected based on areaStat values (B). The first row of two heatmaps at the bottom presents the reference methylation profile of the tumour and normal cell types. Bulks are ordered by the tumour purity and the original methylation beta-values of the bulks are shown in the top heatmap. **C.** Absolute error between reconstructed and reference methylation levels (purple), and between reconstructed and bulk methylation levels (green) for regions selected based on areaStat. Statistics were calculated using paired t-test with Bonferroni correction.

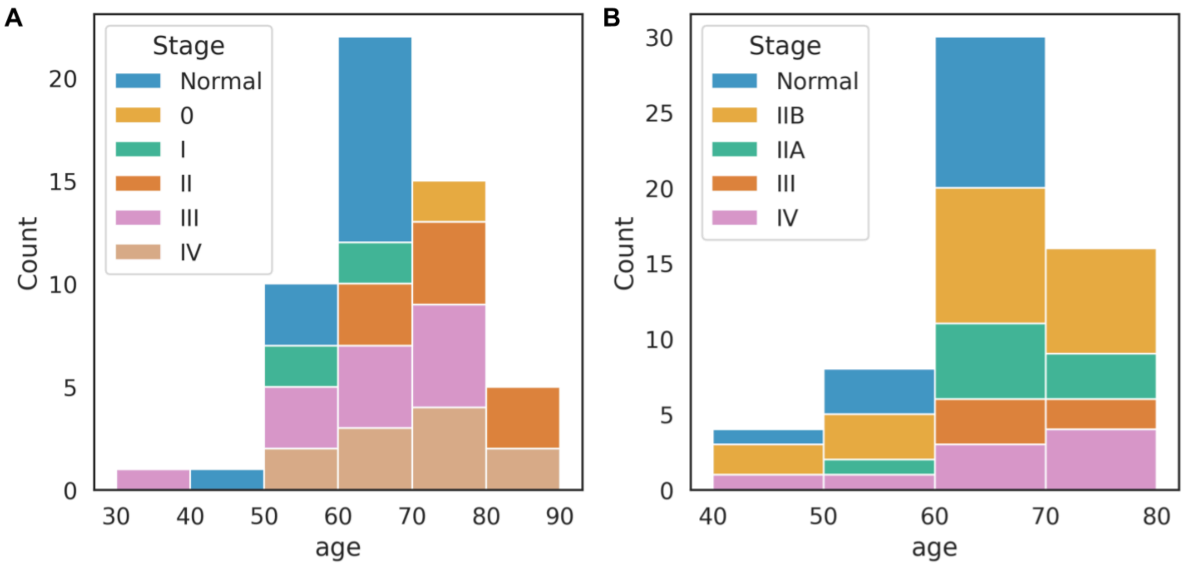


**Supplementary Figure 6.** Age histogram of cancer patients coloured by cancer stages in ctDNA samples from CRC patients (A) and from PDAC patients (B).

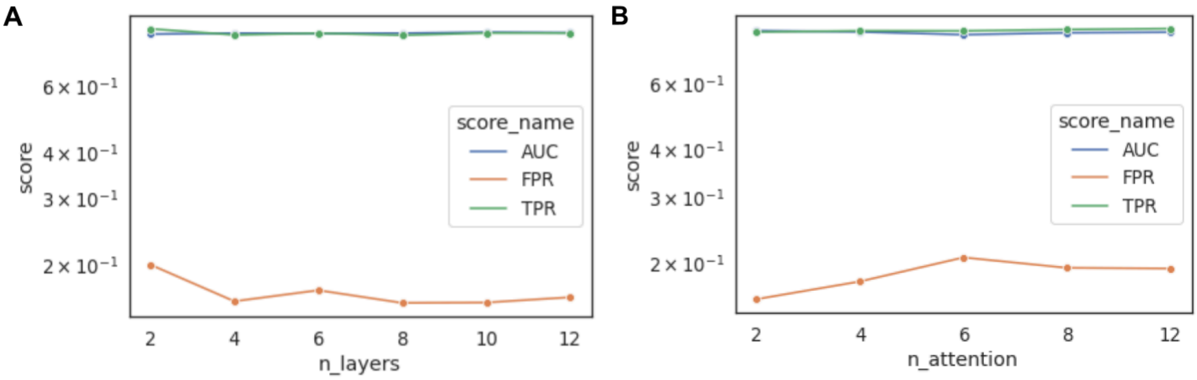


**Supplementary Figure 7.** Comparison of read-level methylome classification performance across different numbers of encoder layers (A) and different numbers of attention heads (B).

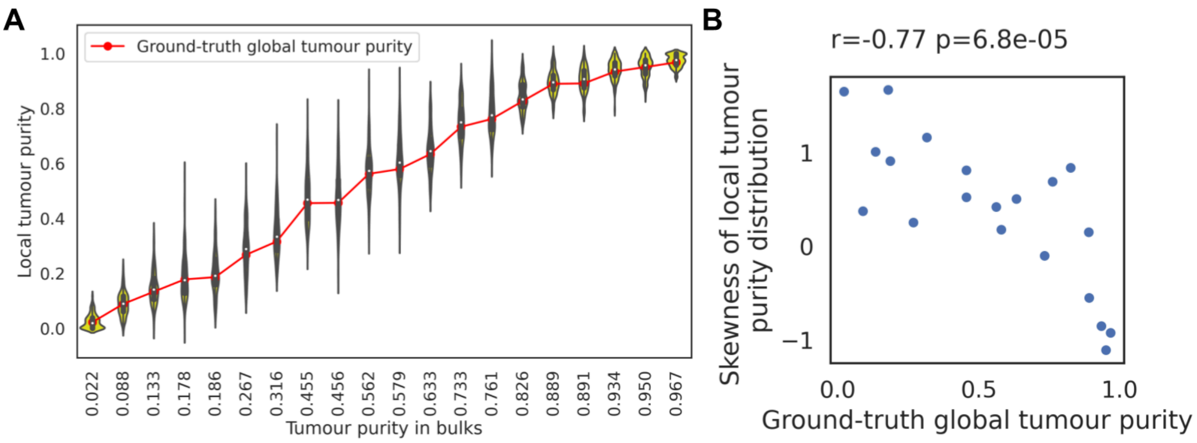


**Supplementary Figure 8.** Region-wise tumour purity in DLBCL pseudo-bulk samples. **A.** Distribution of region-wise tumour purity (local tumour purity) is presented in the violin plot. The ground-truth tumour purity of bulks is shown in the red line. The samples are ordered by the ground-truth value. **B.** Correlation between the skewness of the local tumour purity values and the ground-truth tumour purity.

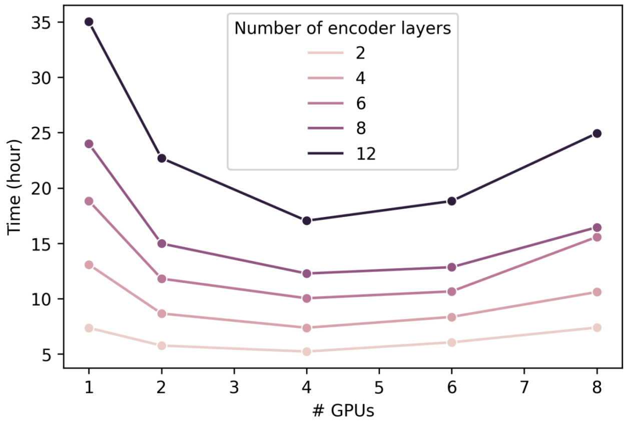


**Supplementary Figure 9.** Running time of MethylBERT by the number of GPUs and the number of encoder layers. Each run was done over 1000 steps.

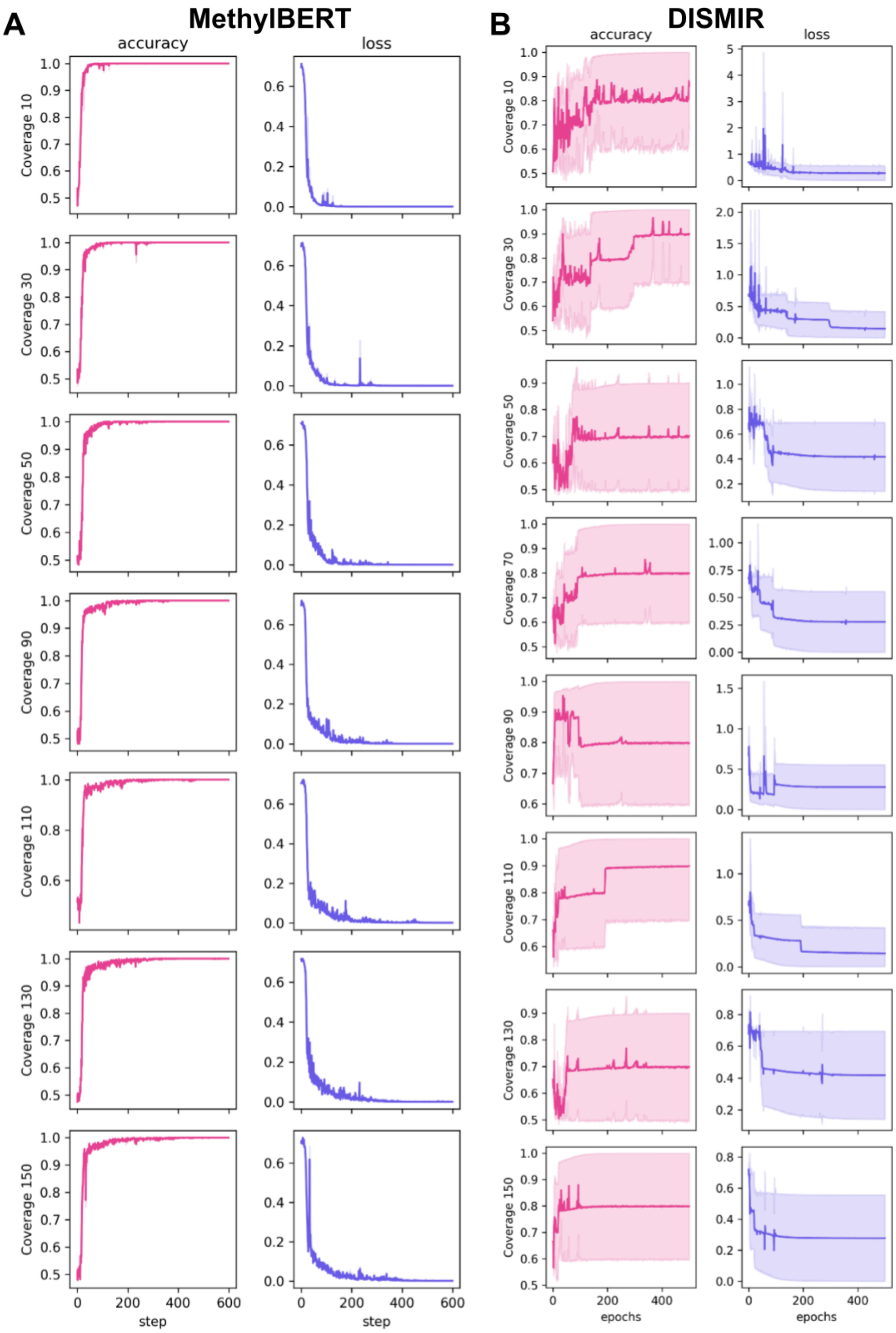


**Supplementary Figure 10.** Read classification accuracy and loss values for different read coverage (in the training set) yielded by MethylBERT (A) and DISMIR (B). We performed five times of the same training for both methods and the line plots show the 95% confidence interval and the mean over the five training runs.

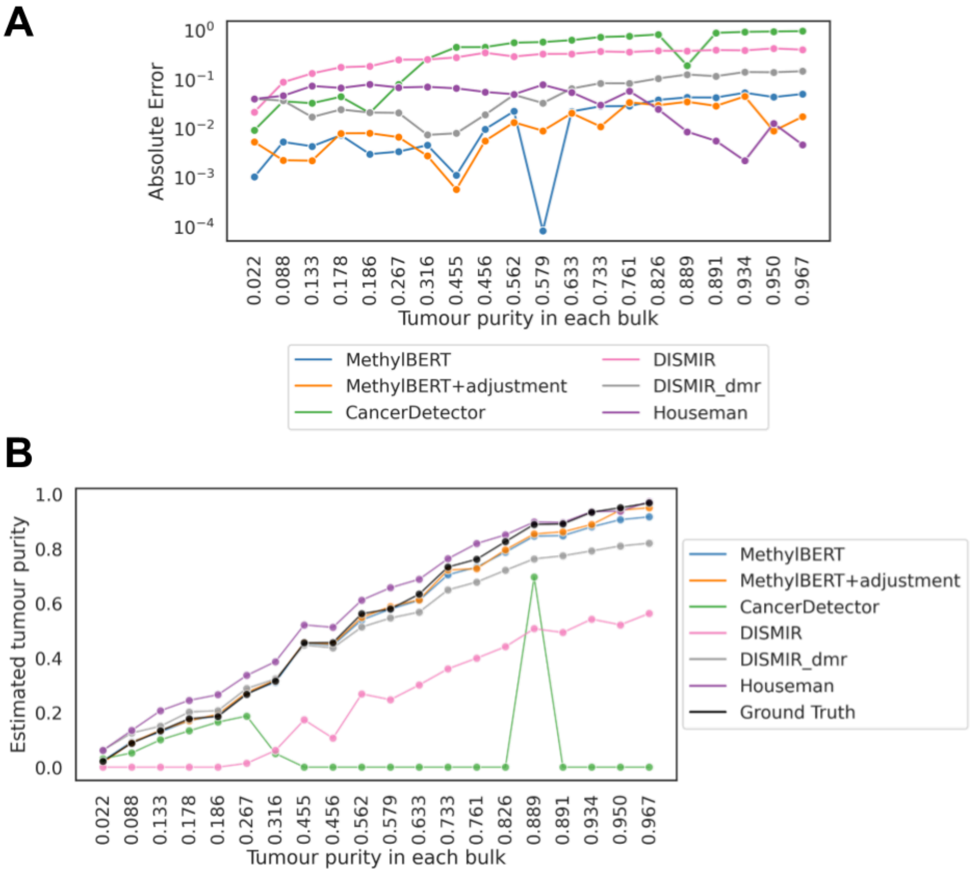


**Supplementary Figure 11.** Tumour purity estimation results for DLBCL pseudo-bulk samples. (A) Absolute error between the ground-truth and estimated tumour purities for each bulk. (B) Estimated tumour purities by individual methods and the ground-truth values for each bulk. Bulks are sorted according to the tumour purity on the x-axis in both plots.

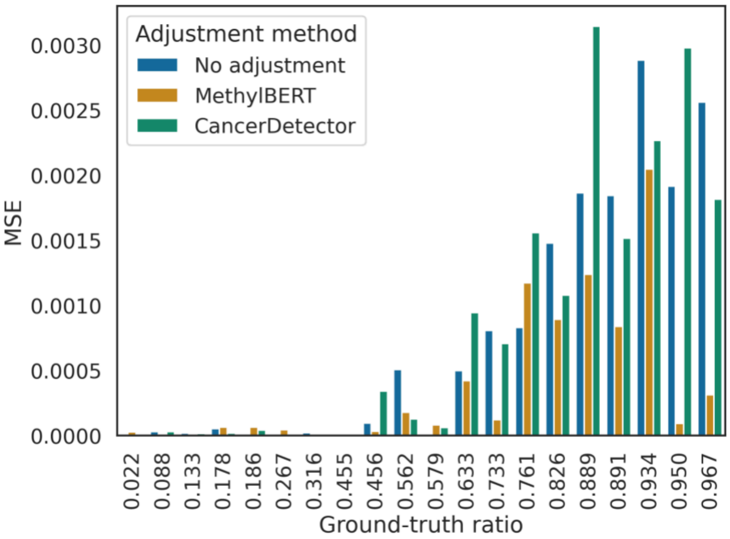


**Supplementary Figure 12.** Performance comparison of different estimation adjustment methods. The histogram shows the mean squared error between the ground-truth and the estimated tumour purity for pseudo-bulks sorted by the ground truth.

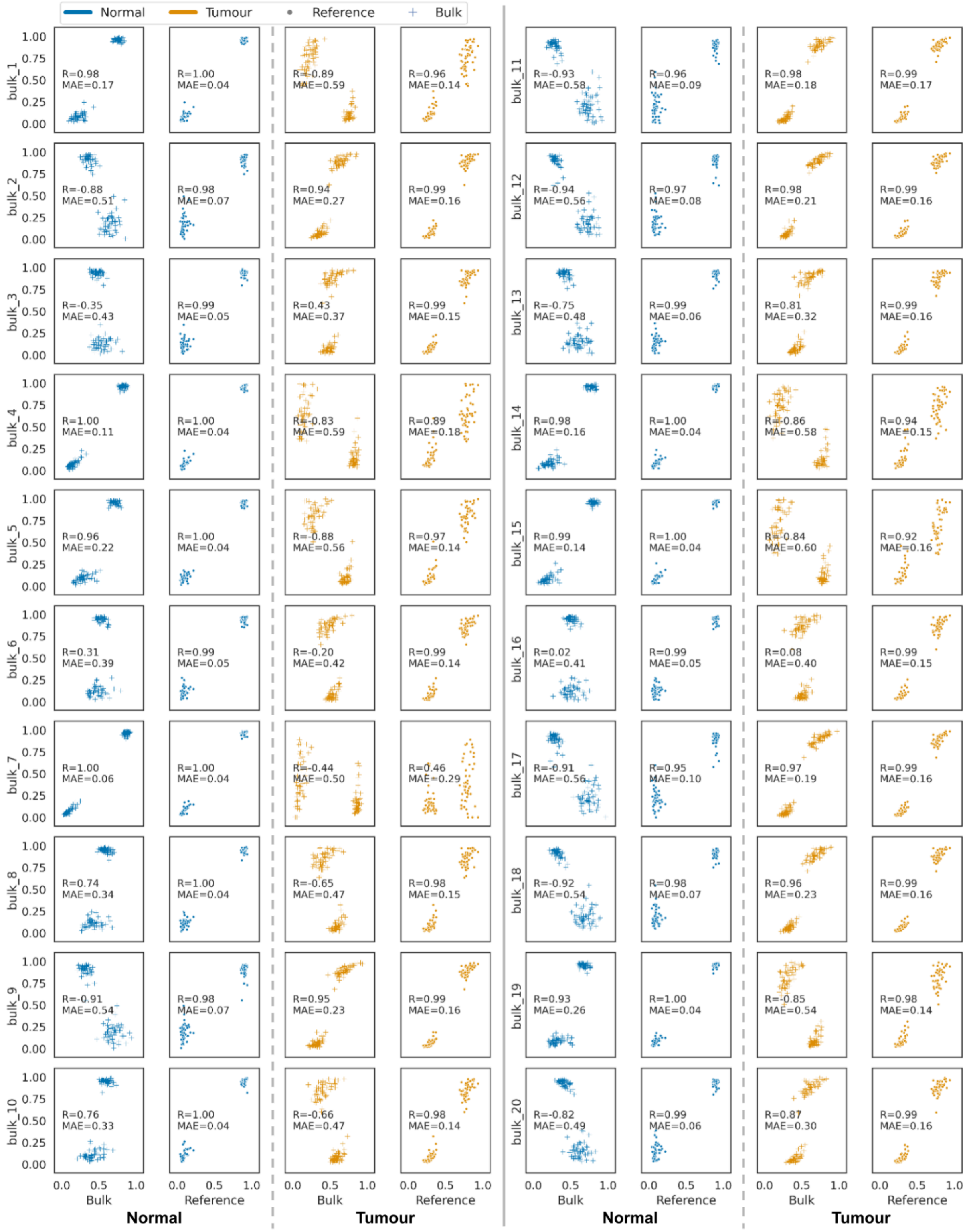


**Supplementary Figure 13.** Reconstructed cell type-specific methylation levels in DMRs (y-axis) compared to the bulk methylation profile (+ marker, x-axis) and the reference cell-type methylation profile (● marker, x-axis). Half tumour-hypermethylated and half tumour-hypomethylated regions were selected. Blue and orange colour indicates the normal and tumour cell types, respectively. In each analysis, mean absolute error (MAE) and Pearson correlation (R) were calculated.

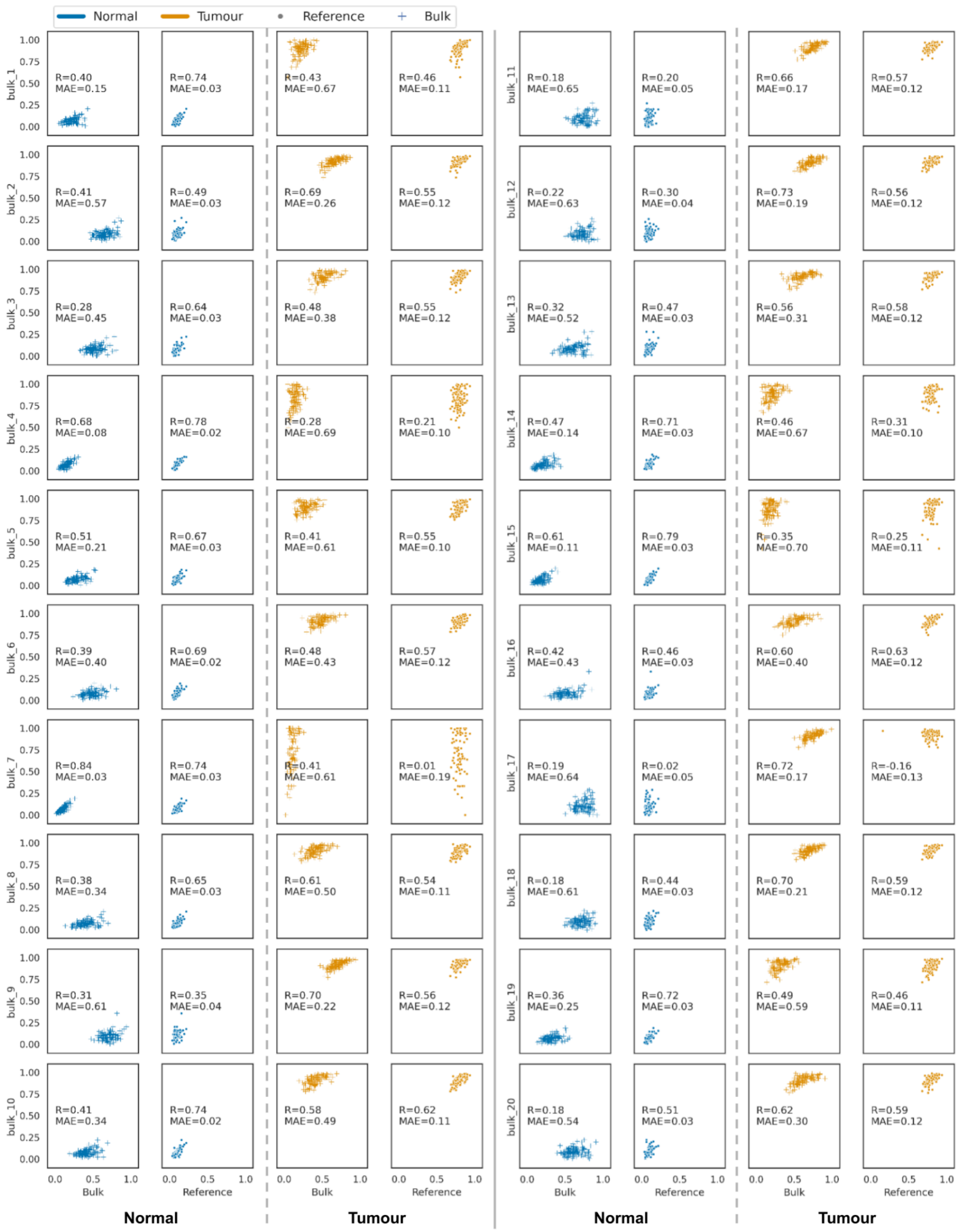

**Supplementary Figure 14.** Reconstructed cell type-specific methylation levels in DMRs (y-axis) compared to the bulk methylation profile (+ marker, x-axis) and the reference cell-type methylation profile (● marker, x-axis). 100 regions with the highest areaStat score were selected. Blue and orange colour indicates the normal and tumour cell types, respectively. In each analysis, mean absolute error (MAE) and Pearson correlation (R) were calculated.
